# Tactile illusion reveals central neural basis for touch pleasantness

**DOI:** 10.1101/2025.03.02.640981

**Authors:** Laura J Pehkonen, Ilona Szczot, Håkan Olausson, Sarah McIntyre

## Abstract

C-low threshold mechanoreceptors (C-LTMRs) have been highlighted in mediating the pleasantness of slow, caressing touch. This has given rise to the “Affective Touch Hypothesis,” which posits that velocity tuning of C-LTMRs underpins pleasantness perception. The known activation preferences of C-LTMRs have been used as a proxy for pleasant touch, yet recent findings have raised questions about the necessity of this peripheral mechanism.

This project explored the peripheral and central mechanisms in affective touch through comparing two motion-conditions: gentle brushing-like motion and apparent motion, an illusory perception of movement produced by successive touches along the skin. We used this illusion to examine whether previously established velocity tuning of true lateral motion is observed in apparent motion, when local information provided to individual peripheral afferents is held constant.

The sole dependence on peripheral modulation predicts that the characteristic inverted-U-shaped relationship between velocity and pleasantness, regularly associated with C-LTMRs, would only be observed for brushing-like motion. Central modulation would instead predict a more similar relationship between the motion-conditions. To investigate this, pleasantness-ratings and velocity-ratings were collected across different velocities (0.1-30cm/s, N=23) for both conditions.

Linear and quadratic regression analyses were performed and for both conditions adding a quadratic term improved the overall model fit, reaching significance (p<0.001). The quadratic term coefficients were negative for both conditions, displaying an inverted-U-shape. Further analyses revealed that motion-condition did not significantly alter the relationship between stimulus-velocity and pleasantness. These findings suggest that the velocity tuning of pleasantness cannot solely be attributed to velocity tuning of individual C-LTMRs.

**Key points:**

- C-low threshold mechanoreceptors (C-LTMRs) have been shown to display a unique inverted-U-shaped relationship between firing frequencies and stroking velocity, which is not seen with any other cutaneous afferents
- An inverted-U-shaped relationship is also observed between perceived touch pleasantness and stroking velocity, giving rise to a hypothesis which posits that the velocity tuning of C-LTMRs underpins the perception of pleasant touch
- We compared traditional stroking (brushing-like motion) to an illusory perception of movement produced by successive touches along the skin (apparent motion) to examine if similar velocity-dependent pleasantness patterns would be observed when information provided to individual peripheral afferents is held constant
- In our experiment, the type of motion did not significantly alter the relationship between perceived pleasantness and stimulus-velocity
- These findings suggest that the velocity tuning of pleasantness cannot solely be attributed to velocity tuning of individual C-LTMRs

## Introduction

The ability to feel and interpret tactile stimulation is crucial, contributing to the dialogue between an individual and their environment in multiple ways, from dexterous manipulation and body awareness to the mediation of social interactions and emotions (Morrison et al. 2010; McGlone et al. 2007). The social and emotional aspects of touch have been interlinked with a specific group of peripheral nerve afferents, C low threshold mechanoreceptors (C-LTMRs, also known as CTs), which have been hypothesized to play a key role in the emotional, hormonal, and behavioural responses to human skin-to-skin contact – forming the basis of the ‘Affective Touch Hypothesis’ (Löken et al. 2009; Olausson et al. 2010). However, emerging evidence suggests that the role of C-LTMRs is far more complex than the “Affective Touch Hypothesis” implies.

### C-LTMRs and Affective touch

Tactile perception has its neural basis in the somatosensory system, which encompasses peripheral nerve afferents, specialized receptors, and the central and spinal components that mediate both proprioceptive and cutaneous processing. In addition to touch, the cutaneous inputs received by the somatosensory system can convey temperature, itch, and pain.

Historically, touch sensation in the human peripheral nervous system was thought to be mediated exclusively by large, myelinated afferents, the A-beta fibres. However, a complementary tactile system was discovered by Zotterman over 80 years ago in experiments utilising cats (Zotterman 1939). This system comprised unmyelinated C-fibers which responded vigorously to light, innocuous touch, in contrast to their nociceptive relatives.

Later, in microneurography recordings, these unmyelinated C-LTMRs have been found in humans, first from the infraorbital and supraorbital nerves and soon after in the arm and leg (Johansson et al. 1988; Nordin 1990; Vallbo et al. 1999), suggesting a more general presence in the hairy skin of humans. Isolated observations of C-LTMRs have recently been reported in the glabrous skin of the palm, although it remains uncertain if these have a similar function to those found in hairy skin (Watkins et al. 2022).

Since the existence of C-LTMRs was confirmed on a larger scale, different methods – notably microneurography, psychophysics and functional neuroimaging – have been utilised to map their characteristics and properties (Morrison et al. 2010; McGlone et al. 2007; Olausson et al. 2010; Nordin 1990; Vallbo et al. 1999; McGlone et al. 2014; Perini et al. 2015; Sanchez Panchuelo et al. 2016; Olausson et al. 2002). Knowing these characteristics and properties help us understand how C-LTMRs achieve their function and allow us to predict their responses to different stimuli.

Similar to nociceptive C-fibres, C-LTMRs are unmyelinated and small in diameter and as a result have slower conduction velocities than myelinated A-fibres. This conduction velocity has been measured to vary between 0.6 and 1.3 m/s (Olausson et al. 2010; Vallbo et al. 1999; Löken et al. 2009). In receptive field properties, C-LTMRs exhibit high heterogeneity. Generally, the receptive field consists of highly sensitive spots, irregularly distributed over a roughly oval shaped area. The number of sensitive spots varies between one and nine (median: 3.8).

Similarly, the area of the receptive field varies between C-LTMRs and is influenced by the force applied locally (mean area: 14.7 mm^2^, SD = 10.5 mm^2^) (Wessberg et al. 2003). Additionally, C-LTMRs have been shown to form an anatomical relationship of functional relevance to hair follicles in the human hairy skin, responding to hair deflection similarly to gentle brushing (Moore et al., 2025).

The functional role of C-LTMRs had been intriguing researchers within the field of neurophysiology since their presence was confirmed in the human hairy skin with early theories centring around a connection with limbic function and skin-to-skin contact (Vallbo et al. 1999; Olausson et al. 2002; 2008). It was after two decades and the findings of Löken et al. in 2009 that a leading theory on this functional role emerged. In their study combining microneurography and psychophysics, Löken and colleagues observed a peculiar and unique relationship between the firing frequency of C-LTMRs, subjective pleasantness-ratings and the velocity used to gently brush the skin. Both the mean C-LTMRs firing frequencies and pleasantness-ratings followed a unique inverted-U shaped path when plotted with the logarithm (log10) of velocity. This relationship in turn showed a significant correlation between the firing frequencies and pleasantness ratings. For other types of LTMRs, the relationship between firing frequencies and velocity was monotonic (Löken et al. 2009).

The characteristics of C-LTMRs have been further studied under different experimental conditions, including chemical agents, locations, textures, and temperatures (Ackerley et al. 2014). Some of the subsequent findings have been utilized to build and support the “affective touch hypothesis”. This hypothesis suggests that C-LTMRs, due to their unique properties, play an important and particular role in the emotional, hormonal (Walker et al. 2017; Portnova et al. 2020), and behavioural (Perini et al. 2015) responses to inter-human, skin-to-skin contact – a tactile construct which can be referred to as affective touch. When experimenting on different temperatures (Ackerley et al. 2014), participants rated all velocities from 0.3 cm/s to 30 cm/s as consistently more pleasant at a neutral – skin-like – temperature (32 °C) compared with a cooler (18 °C) and warmer (42 °C) temperatures. In an experiment by Croy et al. (2016), all participants were observed stroking their partner within the velocity-range (range: 1-10 cm/s) which elicits the highest firing frequencies in C-LTMRs, often referred to as “CT-optimal touch”. Additionally, gentle brushing has been rated as significantly less pleasant by patients with hereditary sensory and autonomic neuropathy type V (HSAN V, a condition characterised by a reduced density of C-fibres and Aδ-fibres) when compared to healthy controls (Morrison et al. 2011), suggesting a correlation between pleasantness and the density of C-LTMR afferents.

With these findings and the characteristics of C-LTMRs in mind, it has become easier to oversimplify the affective touch hypothesis and assume pleasant touch as a proxy for C-LTMR activity. There, however, exists evidence sprinkled in thorough the years suggesting a more complicated story behind the function and purpose of C-LTMRs. The issue of over-interpretation was touched upon by Croy et al. (2021) in a review of five earlier pleasantness experiments.

Although a quadratic fit on the group level was highly replicable, large individual variabilities exist. In the review, only 42% of the individual participants produced a negative quadratic fit (corresponding to the characteristic inverted U-shaped curve) between pleasantness and velocity.

In several prior studies involving “*CT-optimal touch*”, the glabrous skin of the palm has been used as a control-location, based on the notion that glabrous skin is not innervated by C-LTMRs and should therefore elicit different patterns of velocity-dependent pleasantness. However, when 18 such studies were included in a meta-analysis, no significant preference for affective touch was observed, when comparing the palm and the forearm (Cruciani et al. 2021). A high correlation between pleasantness ratings in the hairy and the glabrous skin was also observed in the review by Croy et al. (2021). Recently, isolated observations of C-LTMRs in the glabrous skin of the palm have been reported, although their sparsity leaves doubt as to whether these can have a similar function to those found in hairy skin (Watkins et al. 2022; Watkins et al. 2021).

Patterns in pleasantness have also been investigated in the hairy skin under experimental conditions eliminating either C-fibre or A-fibre function. In a study by Marshall et al. (2019), pleasantness-ratings were collected from patients undergoing anterolateral cervical cordotomy, before and after the intervention. The goal of this intervention was to alleviate cancer-related pain through lesioning of the spinothalamic tract, which is thought to contain the spinal projections for CT-afferents (Olausson et al. 2010; Abraira and Ginty 2013). However, no significant main effects of side (cordotomy-control) nor cordotomy status (precordotomy-postcordotomy) on velocity-dependent pleasantness were observed. These findings are incompatible with a model where privileged and anatomically separate second order projections of CT-afferents underpin pleasant touch perception.

Recent results also suggest a role for Aβ afferents in processing affective and social touch. Significant changes in pleasantness were observed in the presence of two types of nerve block, selectively targeting and reducing A-fibre function. Both forms of nerve block, ischemic and non-ischemic compression block, led to a nearly complete loss of pleasantness for brushing, suggesting that the contribution of A-fibre afferents is necessary for pleasantness-perception in typically developed individuals (Case et al. 2023). To further support potential A-fibre contribution in affective touch, microneurography recordings have shown that Aβ-fibres can effectively encode and discriminate different social touch expressions (Xu et al. 2024).

Additionally, possible evidence suggesting top-down modulation of pleasantness-perception has been obtained in experiments utilising different odours. In a study by Croy et al. (2014), unpleasant odour significantly decreased the perceived pleasantness of touch at two stroking velocities (3 cm/s, 30 cm/s). The magnitude of this modulation was further influenced by individual disgust sensitivity. Similarly, a subsequent study observed that disgust towards a given odour was negatively correlated with touch pleasantness, but not intensity (Croy et al. 2016).

### Using illusory movement to manipulate peripheral neural activation

The objective of this project was to further explore the peripheral mechanisms of affective touch and pleasantness-perception utilising apparent motion. Apparent motion is an illusory perception of movement, achieved through the presentation of individual stimuli in sequence (Ekroll, Faul, and Golz 2008; Steinman and Pizlo 2000). A familiar visual analogue of this illusion is found in the function of modern television screens and computer monitors in which the activation of successive pixels results in the illusion of movement across the screen.

In a tactile context, this illusion of movement arises when two or more separate tactile stimuli (e.g. indentation or vibration) are presented successively on the skin surface, at different locations along a trajectory (Kirman 1974; Kirman 1983). When successfully executed, this stimulation pattern is perceived as continuous motion along the area encompassed by all the stimulation points, even though the individual actuators do not move across the surface of the skin (Hachisu and Suzuki 2019; Kuroki et al. 2012; McIntyre et al. 2014; McIntyre et al. 2016).

Since individual actuators only contact the skin at one small area, if the distances between contact points is greater than the receptive field sizes of primary afferents then each one activates a different set of primary afferents. The perception of movement in the apparent motion illusion must therefore arise from central processing that integrates signals from these different sets of primary afferent inputs. At the same time as the illusory global motion and its properties are perceived (e.g. direction and speed), so can the local mechanical inputs be perceived at the individual contact sites (e.g. speed and force of indentation).

By using apparent motion in this study, the goal is to hold constant any local contact cues – including differences in contact time, force, or lateral motion – that could influence the response of any individual C-LTMRs, while still being able to manipulate the global velocity of apparent motion (Fig. 1). The perception of apparent motion relies on the integration of tactile input across the different locations on the skin, a process shaped by the organization of receptive fields. These fields exist at both the peripheral and cortical levels, enabling the nervous system to process touch information with varying degrees of spatial resolution and integration.

**Figure 1.**
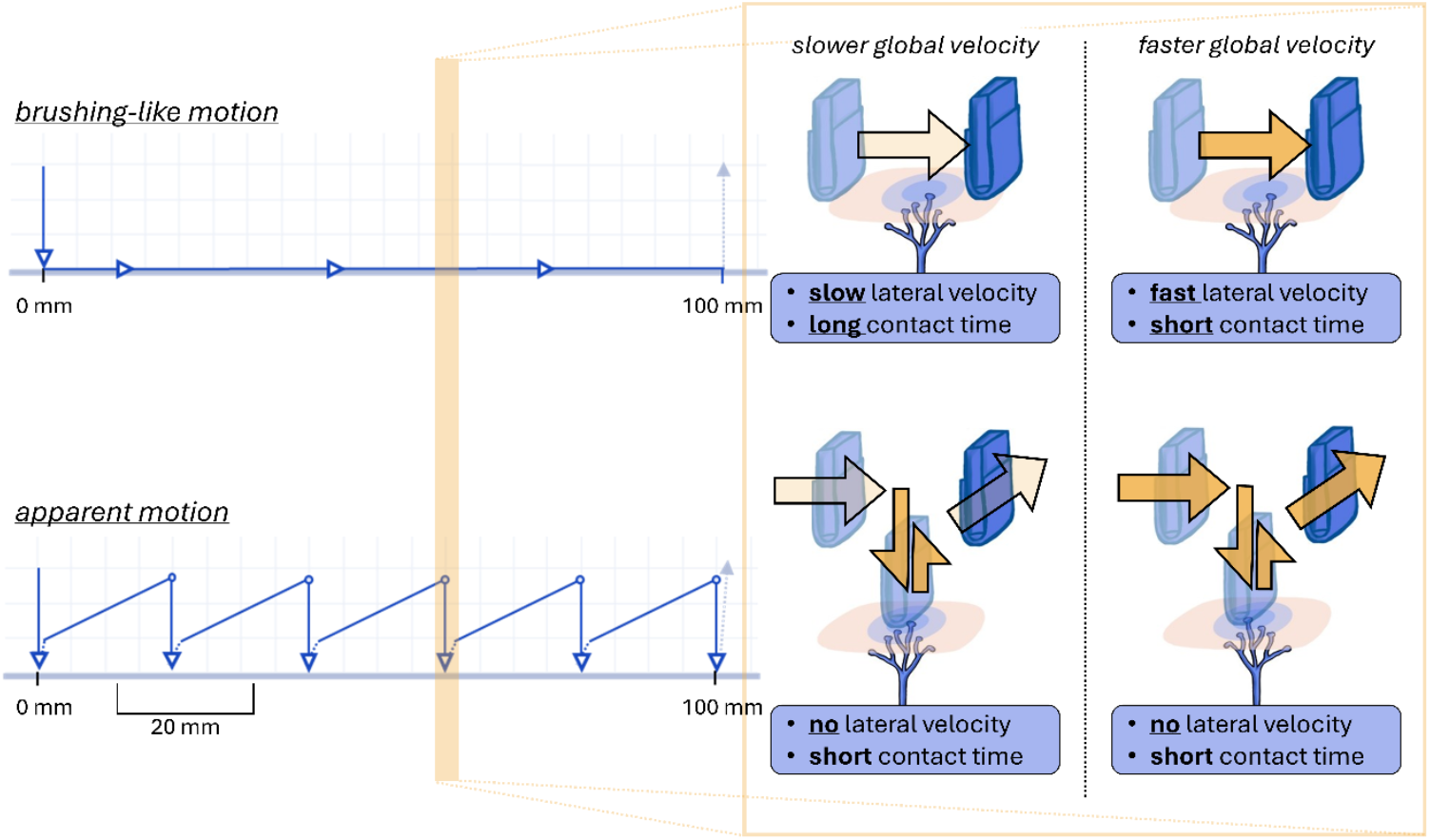
Diagrams of the brushing-like motion and apparent motion trajectories. On the right, a simplification of the methodological goal from the perspective of an individual CT-afferent. When subjected to brushing-like motion, a single CT-afferent can tune its response depending on velocity (Löken et al. 200S). With apparent motion, where all contact-characteristics are kept similar across global velocities, an individual CT-afferent cannot provide velocity-dependent response.

On the peripheral level, each tactile afferent has a localized and spatially restricted skin region where the mechanoreceptors respond to stimuli. These peripheral receptive fields (PRFs) vary in size and density across different sites on the body, influencing tactile acuity (Johansson and Vallbo 1979; Corniani and Saal 2020; Abraira and Ginty 2013).

As multiple PRFs converge in the somatosensory cortex, cortical receptive fields (CRFs) emerge, building a hierarchical network (Sanchez Panchuelo et al. 2016). In the primary somatosensory cortex, neurons maintain a somatotopically organized map, with receptive fields increasing in size when moving from the anterior to the posterior. Higher-order areas, such as the secondary somatosensory cortex S2, present with even larger receptive fields, enabling increasing levels of integration (Pei and Bensmaia 2014; Sanchez Panchuelo et al. 2016; Pei et al. 2010).

It is at these cortical levels where the sequential tactile stimuli of apparent motion are interpreted as continuous motion with a specific global velocity. In the brushing-like motion condition, the peripheral neural response is tuned to velocity – the local, lateral velocity across the peripheral receptive fields – due to the specific velocity-dependent firing rates of the peripheral mechanoreceptors. In contrast with apparent motion, each of the contacts on the peripheral level are held constant. Only the interstimulus – time between two subsequent stimuli – intervals are varied to control global velocity. It is the integration of these distinct sequential stimuli from different locations on the cortical level which creates the perception of motion.

As for this experiment, two alternatives were seen as the possible outcome: apparent motion and brushing motion yield either similar or differing patterns regarding pleasantness and velocity. If C-LTMRs play a truly particular role in pleasantness perception, apparent motion should not be able to recreate the negative quadratic relationship between pleasantness and velocity. This would be a result of the peripheral C-LTMRs not being able to encode velocity in their firing rates. If, however, a negative quadratic relationship would be observed in apparent motion, it would suggest that the peripheral velocity tuning of C-LTMRs cannot solely explain velocity dependent pleasantness.

## Methods

In this study, the effects of two different tactile stimuli at an array of global velocities were investigated on perceived pleasantness. The tactile stimuli were delivered using a robotic arm with a soft silicon contactor. The two stimuli were apparent motion, and a brushing-like motion similar to previous studies on affective touch. Pleasantness ratings and perceived velocity were collected using a Visual Analogue Scale (VAS). A validation experiment using a force sensor and video camera was performed to confirm the force and time characteristics of the stimuli.

### Participants

The experiment was conducted in accordance with the Declaration of Helsinki and GDPR guidelines and received approval from the Swedish Ethical Review Authority (dnr 2022-01108-02). Informed, written, consent was obtained before starting the experiment. The experiment included a total of 23 healthy volunteers (13 females, 10 males), with a median age of 24 (range 20-58).

### Equipment

The tactile stimuli were administered using the Universal Robots UR3e cobot (Odense, Denmark), equipped with a custom silicone contactor (See Supplementary Materials, Figure 1). The contactor was designed to imitate a soft brush, popular in prior experiments; the edge in contact with the skin had a width of 5 cm and thickness of 1 cm. A thin layer of talc-powder (Johnson & Johnson, New Brunswick, United States) was applied on the contactor surface to reduce friction and prevent the skin and hair from sticking. A force-torque sensor (Nordbo Robotics, Odense, Denmark) was placed between the robotic arm and the contactor to monitor and validate applied force. The same equipment was used for both tactile conditions.

### Stimulus design

For both the brushing-like motion and apparent motion, six different global velocities (0.1, 0.3, 1, 3, 10 and 30 cm/s) were selected, similarly to the velocities used in the experiment by Löken et al. (2009). The stimuli were delivered with the gentle force of 0.4 Newtons (mean 0.415 N, SD: 0.0407 N), across a trajectory of a 100 mm on the left dorsal forearm, starting 1 cm proximal to the wrist. A diagram showing apparent motion and brushing-like motion is shown in Figure 1.

#### Control condition, brushing-like motion

Continuous stroking motion was applied along the entire length of the trajectory (10 cm) on the forearm at a constant force, using the six velocities. For this condition, the global velocity was the same as the local lateral velocity at skin-contact. The duration of skin-contact depended on this stimulus-velocity (t_contact_ = 10 cm / v). The brushing-like stimuli resemble stroking movements previously shown to activate C-LTMRs

#### Apparent motion condition

Sequential taps were administered at six distinct locations along the trajectory with a 20 mm distance between the centre of each tap. With the contactor thickness of 10 mm, a 10 mm long patch of skin received no tactile stimulation in between the taps. The contact time and force with each tap remained constant, regardless of global velocity. Only the interstimulus interval was varied to modulate global velocity. These taps are likely to activate C-LTMRs (as well as A-beta afferents) as they resemble indenting stimuli that can activate C-LTMRs to a similar degree to stroking movements (Middleton et al. 2022; Xu et al. 2024). However, since the local information at skin-contact is held constant, the individual peripheral neurons cannot tune their response to the global velocity of the stimulus.

### The experimental setup

Since both the brushing-like motion and apparent motion utilised the same equipment, randomization of the stimuli across conditions was possible. Each unique stimulus (condition, velocity) was presented three times in a random order, with the entire experiment protocol consisting of 36 stimuli. In addition to the participants response time, each stimulus was preceded by a pseudorandom pause between 6 and 15 seconds (t_pause_ = 5 + randint(1, 10) s). Before the start of the protocol, a pre-defined set of stimuli was presented to validate the arm position and to introduce the participant to the stimuli.

During the experiment, the participant was seated in a comfortable chair. The participant’s left arm was placed on an armrest with the dorsal forearm facing upwards. This area was visually isolated from the participant with a sheet so they could not see the movements of the cobot arm (Figure 2). Additionally, the participant wore noise-cancelling headphones playing white noise for auditory isolation so they could not hear the motors in the cobot arm. All experiments were conducted with the same set-up and by the same investigator.

**Figure 2.**
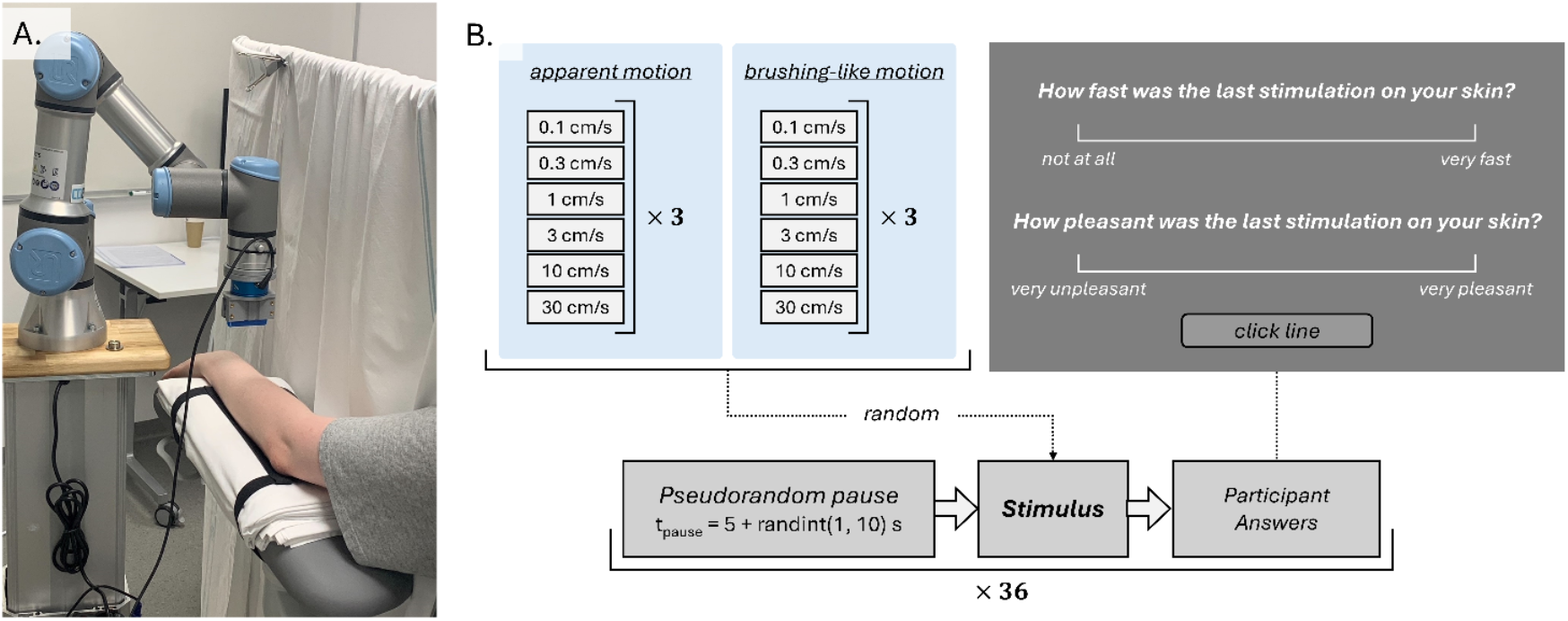
A. and B. show the experimental layout from two different perspectives. Participants placed their left forearm on an armrest, where all tactile stimulation was provided using the UR3e cobot. This was visually separated from the participant with a sheet. For auditory isolation, participants wore noise-cancelling headphones playing white noise. The right arm was used to answer questions on a laptop. C. shows a schematic over the experimental protocol. A total of 12 unique stimuli were presented three times in a random order, preceded by a pseudorandom pause between 6 and 15 seconds. Participants would then provide their answer using a VAS.

### Data-acquisition and analysis

After each stimulus, the participants were presented with two questions:

- “How fast was the last stimulus on your skin?”
- “How pleasant was the last stimulation on your skin?”

In addition to pleasantness, data on the subjective perception of velocity was collected. This measurement was added to assess the perception of global velocity between the two tactile conditions. Additionally, it allows investigation of the relationship between perceived velocity, global velocity, and pleasantness.

For data collection, the Visual Analogue Scale (VAS) was used (Figure 2c). VAS is a psychometric tool consisting of a continuous scale. Anchor-labels are used to define the scale, with no numerical marks presented to the answering party. An answer is given by marking a spot along the scale which is later transformed into an arbitrary figure for data-analysis. For the experiment, anchor-labels were assigned to the extremes, which were given the numeric values of -10 and 10. The extremes for perceived velocity were “not at all” and “very fast”, and for pleasantness were “very unpleasant” and “very pleasant”.

Descriptive statistics and data-analysis were performed using SPSS (IBM, version 29.0.0). Before conducting regression analyses, velocity-categories were transformed to log10 values, consistent with previous studies.

To assess the relationship between velocity and pleasantness across the two tactile conditions, regression analyses were conducted using both linear and quadratic models. To account for inter-individual variability and repeated measurements within participants, multilevel mixed-model analyses were employed. This approach was chosen because it allows for the inclusion of both fixed effects (stimulus type and velocity) and random effects (participant-level differences), providing a more robust and generalized analysis. Maximum likelihood estimation was used to ensure optimal model fit, while a random intercepts model controlled for baseline differences between individuals.

Additionally, to explore how subjective velocity ratings corresponded to global stimulus velocity and pleasantness, Spearman’s rho correlation, curve fitting, and multilevel mixed-model analyses were performed.

### Validation experiment

A validation experiment was performed to evaluate the contact-time in the apparent motion stimuli-set. Two measurement methods were used: real-time force recordings from the force-torque sensor, sampled at 240 Hz, and video recordings captured at 240 frames per second. During a session, apparent motion programs were recorded simultaneously using video and real-time force data. Data collection was performed for 6, 9, and 15 repetitions for the velocities of 0.1 cm/s, 0.3 cm/s, and 1–30 cm/s, respectively. For force recordings, the duration of sustained downward force corresponding to a tap was extracted. In the video recordings, individual frames where contact and/or skin displacement could be observed were manually counted to estimate contact time.

No statistically significant trends in contact times across velocities were found using the Jonckheere-Terpstra Test for Ordered Alternatives (p = 0.0502 for video frames, p = 0.327 for force recordings). This test was chosen due to its suitability for detecting ordered differences in a non-parametric dataset across multiple groups, making it appropriate for evaluating trends in contact times across predefined velocity conditions.

Contact times derived from frame counting were consistently higher (median: 0.0792 s, IQR: 0.0083 s) compared to those obtained from force recordings (median: 0.0640 s, IQR: 0.0120 s). However, the difference between the two methods did not significantly vary across velocities (Jonckheere-Terpstra p = 0.582). When averaging the two methods, the median contact time was 0.0718 s (IQR: 0.0082 s).

Additionally, the real-time force recordings revealed that the global velocities for apparent motion stimuli did not reach the two highest programmed speeds (10 cm/s and 30 cm/s). Instead, the actual recorded velocities were 5.7 cm/s and 6.8 cm/s, respectively, indicating a deviation from the expected values. For brushing-like motion stimuli, global velocities followed their respective encoded velocities.

Because of the limitations in achieving the faster sub-optimal velocities in the apparent motion condition, we corrected the velocity groups accordingly for all analyses. The decision to continue was based on the potential for meaningful trends to emerge even within the available range. While 3 cm/s is commonly referenced as the “most” CT-optimal velocity, in previous studies with detailed regression analyses the theoretical peak pleasantness varies – sometimes even reached at velocities around or below 1 cm/s. For example, Schirmer et al. (2022) reported a theoretical peak at 1.025 cm/s, reinforcing the idea that pleasantness trends could still be effectively captured even with a more limited range of velocity.

## Results

### Group-level Linear and Quadratic Regression Analyses

For the two types of motion stimuli, linear and quadratic regression models were performed to evaluate how stimulus-velocity influenced the pleasantness-ratings. While linear regression models did not reveal significant effects of velocity on pleasantness, adding a quadratic term significantly improved the model fit for both conditions.

The simple linear regression model for brushing-like motion was not statistically significant revealing no significant effect of velocity on pleasantness (R = 0.008, R^2^ = 5.63×10^-5^, F(1, 404) = 0.023, p = 0.880, Table 1, Supplementary Table 1). When the quadratic term was added, the model fit improved and reached high statistical significance (R = 0.355, R^2^ = 0.126, F (2, 403) = 29.033, B = -2.44, p = 1.66×10^-12^, Figure 3b, Table 1, Supplementary Table 1).

**Table 1.** Model Summary of the linear and quadratic regression analyses for both apparent motion and brushing-like motion. Dependent variable pleasantness.

| Model | Condition | R | R<sup>2</sup> | SE | F | Sig. |
| --- | --- | --- | --- | --- | --- | --- |
| Linear<sup>1</sup> | Apparent motion | 0.093 | 0.009 | 3.778 | 3.528 | 0.0611 |
|  | Brushing-like motion | 0.008 | 5.63×10<sup>-5</sup> | 4.212 | 0.023 | 0.880 |
| Quadratic<sup>2</sup> | Apparent motion | 0.271 | 0.073 | 3.657 | 16.069 | 1.92×10<sup>-7</sup> |
|  | Brushing-like motion | 0.355 | 0.126 | 3.943 | 29.033 | 1.66×10<sup>-12</sup> |

<sup>1</sup> Predictors: (Constant), $\log_{10}(\text{velocity})$ <sup>2</sup> Predictors: (Constant), $\log_{10}(\text{velocity})$ , $\log_{10}(\text{velocity})^2$

**Figure 3.**
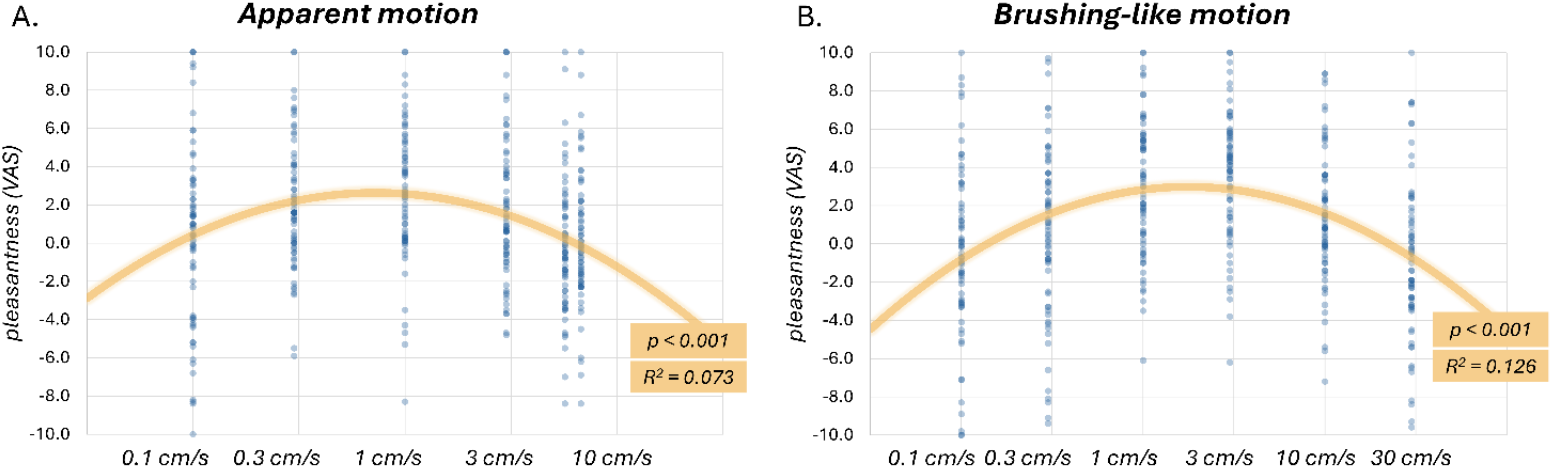
Scatterplot for pleasantness-ratings (y-axis), plot ‘A.’ for apparent motion and ‘B.’ for brushing-like motion. X-axis shows the value of log_10_(velocity), with labels showing the corresponding stimulus velocity. In orange, the best fitting quadratic regression model is presented, including R^2^ and p-value.

Similarly, the linear regression model for apparent motion was not statistically significant (R = 0.093, R^2^ = 0.009, F(1, 407) = 3.528, p = 0.0611, Table 1, Supplementary Table 1). The inclusion of a quadratic term revealed a fit of statistical significance (R = 0.271, R^2^ = 0.073, F(2, 406) = 16.069, B = -2.98, p = 1.92×10^-7^, Figure 3a, Table 1, Supplementary Table 1).

These analyses indicate that pleasantness ratings did not follow a linear relationship with velocity for either brushing-like or apparent motion. However, when a quadratic term was included, the model fit improved significantly for both conditions, suggesting that pleasantness follows an inverted-U function with velocity.

### Multilevel Mixed-Effects Model Analyses

In order to directly compare the relationship between velocity and pleasantness under the two conditions, multilevel mixed-effects models were utilised.

In the multilevel mixed-effects model, a significant main effect of stimulus-type was observed (F (1, 815) = 4.224, p = 0.0402), with apparent motion associated with higher pleasantness-ratings (estimate = 1.238). Both the linear (log_10_velocity, F (1, 815) = 65.646, estimate = 6.088, p = 5.13×10^-14^) and the quadratic (log_10_(velocity)^2^, F (1, 815) = 68.245, estimate = -2.441, p = 7.20×10^-15^) terms for velocity were highly statistically significant (Supplementary Table 2). Neither one of the interaction-terms with velocity, type* log_10_velocity (p = 0.490) nor type* log_10_(velocity)^2^ (p = 0.412), were statistically significant. The absence of significant interaction effects suggests that the velocity-pleasantness relationship follows a similar pattern in both conditions.

### Individual-level Linear and Quadratic Regression Analyses

Linear and quadratic regressions were additionally performed on the individual level. Out of the 23 participants, nine showed a significant linear fit with apparent motion compared to six with brushing-like motion. For the quadratic fit, 12 individuals had a significant quadratic fit for both the apparent motion and brushing-like motion (Supplementary Figures 2, 3, 4 & 6). The *R*-, *R*^*2*^- and *F*-values from these regressions were collected and then compared between the two conditions using related-samples Wilcoxon signed rank test. No statistically significant differences were observed in these values between the two conditions, neither for the linear nor the quadratic regressions (Supplementary Figure 5 & Table 3). These results further suggest that velocity-dependent pleasantness perception is similar between the brushing-like and apparent motion.

**Figure 4.**
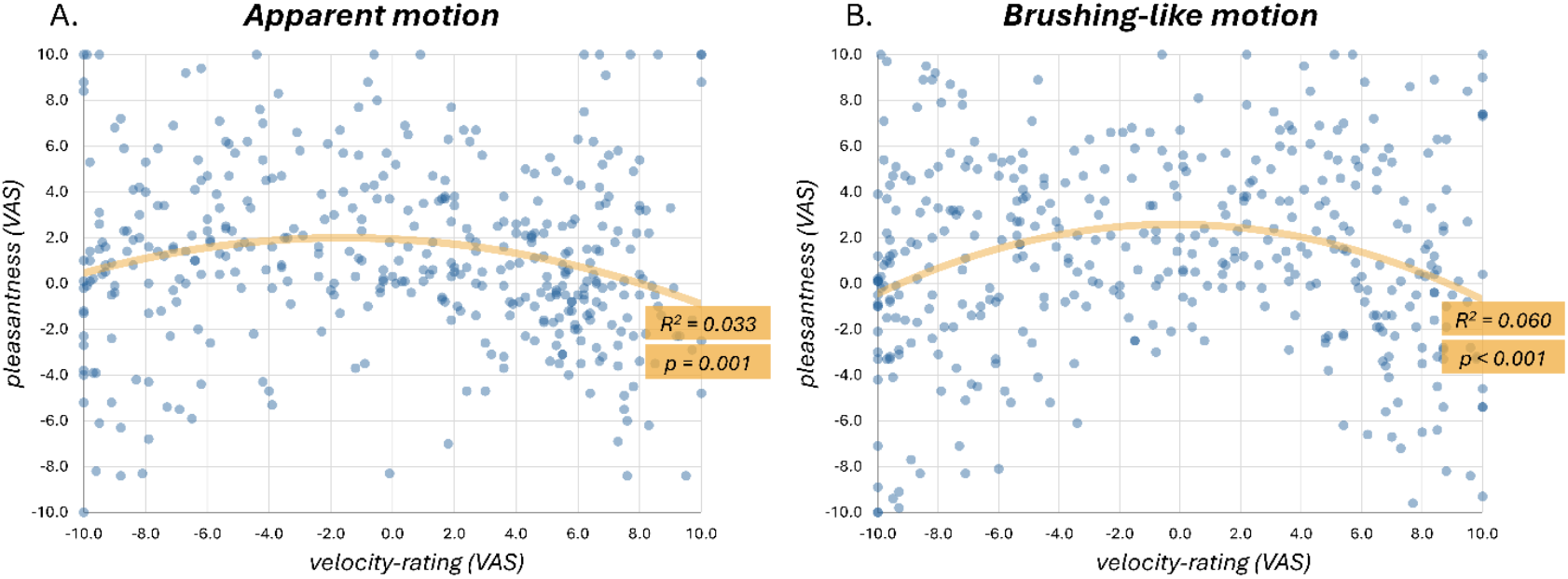
Scatter-plot for pleasantness-ratings (y-axis) and subjective velocity-ratings (x-axis). Plot ‘A.’ shows apparent motion and ‘B.’ brushing-like motion. In orange, the best fitting quadratic regression model is presented, including R^2^ and p-value.

### Subjective velocity ratings

Linear regression analyses, exploring the relationship between velocity-ratings and the log_10_-value of stimulus velocity, revealed similarly significant fits (apparent motion: R = 0.881, R^2^ = 0.776, F(1, 406) = 1408.034, p = 1.13×10^-134^ – brushing-like motion: R = 0.870, R^2^ = 0.757, F(1, 402) = 1251.327, p = 1.49×10^-125^, Supplementary Figure 7 & Table 4), indicating that perceived velocity closely follows actual velocity in both conditions.

A multilevel mixed effects model (Supplementary Table 5) was conducted to observe if perceived velocity would alter the trends in velocity-dependent pleasantness. In this model, neither stimulus-type (p = 0.149) nor the linear velocity-rating term (p = 0.671) yielded significant main effects. A statistically significant main effect was, however, observed with the quadratic term for velocity-ratings (velocity-rating^2^, F(1, 812) = 36.081, estimate = -0.032, p = 1.56×10^-7^). Neither of the interaction terms reached significance (p > 0.200), suggesting that not only does the inverted-U function between pleasantness and velocity translate over to perceived velocity, but that this pattern is consistent between brushing-like and apparent motion (Figure 4).

## Discussion

Our findings suggest that the perception of pleasant touch is not solely driven by the velocity tuning of peripheral C-LTMRs but instead emerges from more complex integration processes at higher levels of the somatosensory system. These findings demonstrate similar patterns in velocity-dependent pleasantness perception in both brushing-like motion and apparent motion.

With the methodological aspects of this project in mind, it can be argued that very similar information is provided to the peripheral cutaneous afferents, regardless of global velocity, in the apparent motion condition. In other words, the individual C-LTMRs cannot receive the necessary information for a velocity-tuned neural response, despite changes in global velocity.

Even without this of C-LTMR input in response to velocity, an inverted U-shaped relationship – similar to that observed in the presence of C-LTMR input (brushing-like motion) – was observed in apparent motion. This in turn suggests that the pattern of velocity tuned pleasantness perception is not a direct product of the peripheral C-LTMR input. Since the perception of tactile motion and its velocity are constructed when peripheral receptive fields are converged into larger cortical receptive fields in the somatosensory cortex, central integration processes appear necessary for pleasantness perception.

Our findings are in line with prior studies targeting and eliminating either C-fibre or A-fibre function. In the experiment involving cordotomy patients by Marshall et al. (2019), no significant effects on pleasantness were observed after lesioning the spinothalamic tract, described to contain the spinal projections for C-LTMRs. This finding suggests that either C-LTMR function is not necessary for pleasantness-perception or the C-LTMR projections travel somewhere outside the spinothalamic tract. The proper function of A-fibres has, however, been shown as necessary for pleasantness perception by Case et al. (2023). The involvement of cortical processes has additionally been suggested in prior experiments utilising different odours, with unpleasant odours decreasing perceived pleasantness.

With the velocity-dependent perception of pleasantness unaffected by both the lack of informative C-LTMR input and the lesioning of the spinothalamic tract (Marshall et al. 2019) but significantly disturbed by the elimination of A-fibre function (Case et al. 2023), it could be argued that C-LTMRs are not an essential component of pleasant touch perception. However, this evidence is limited to adult populations and does not contradict the hypothesis that C-LTMRs may play a crucial role in shaping pleasantness perception during neurodevelopment (Croy et al. 2019).

However, a question can still be raised whether C-LTMRs possess uncharted alternate functions. Such potential functions have been speculated thorough the years, from ticklishness (Zotterman 1939) to interoception (Craig 2002).

### The influence of stimulus features

While standardized tactile stimulation methods enhance comparability across studies, they may limit our understanding of the full complexity of touch, especially naturalistic touch. In studies focusing on affective touch, tactile stimulation is most commonly delivered using either a rotatory tactile stimulator (RTS) or manually, typically with a soft brush measuring 4 to 7 cm in width, originally designed for painting or cosmetic purposes. Although this coherence in methodology across studies can be seen positive from a standardization perspective, it might fail to capture some complexities in tactile interactions between individuals (Maallo et al. 2022) and the environment.

In this experiment, we employed a linear motion trajectory parallel to the surface of the skin, applying tactile stimulation with a custom-designed silicone tool. The trajectory of movement, despite it potentially influencing velocity-dependent pleasantness perception, has received relatively little attention in the context of affective touch research.

A two-part study by Schirmer et al. (2023) explored the effects of different motion trajectories – rotary (RTS), linear, and oval – on perceived pleasantness. Only their linear trajectory exhibited consistently significant quadratic effects. In contrast, the RTS-trajectory showed mixed results, and the oval trajectory failed to produce any significant effects (Schirmer, Cham, et al. 2023). These findings, along with our own results showing a significant negative quadratic term for a linear trajectory, highlight how trajectory shape can influence velocity-dependent pleasantness perception.

Notably, in our study, we chose to abandon the use of the RTS and the traditional soft brush early on to better control the spatiotemporal characteristics of the tactile stimulation. Despite these methodological changes, we successfully replicated the velocity-dependent perception of pleasant touch, characterized by a significant negative quadratic regression. A similar result has even been reported with vibration-based apparent motion Le et al. (2024). This replicability could suggest broader applications for capturing and delivering pleasant touch in experimental settings and in the world of haptic technology. In addition, recent research has shown that C-LTMRs are activated not only by the slow gentle stroking commonly thought of as “CT-optimal” touch, but also by a slow firm press of the hand (Xu et al. 2024), by monofilament indentation (Middleton et al. 2022), and by selective deflection of single hairs (Moore et al. 2025), although their responses to vibration remain uncharacterised in humans. Taken together, this supports recent calls within the community to avoid over-simplifying the relationship between pleasant touch and C-LTMR activation (Schirmer et al. 2023).

### Limitations and future directions

Although the study benefits from a strong methodological design, including randomization, it is not without limitations. The primary limitation is the absence of microneurography data, which could more definitively confirm the similarity of peripheral neural responses elicited by the individual taps in our apparent motion stimulus. However, this limitation presents a clear motivation for future investigations.

Additionally, our robot arm was unable to reach the highest velocities of interest in the apparent motion condition. Despite capturing a robust, quadratic relationship between velocity and pleasantness, it is a possibility that other kinds of relationships would appear with the entire velocity-spectrum included.

This experiment captures only one specific construction of apparent motion, with many alternative configurations possible. For instance, a completely different approach could be taken regarding contact time, which in this experiment was held constant and minimal. As a result, the effect of contact time on pleasantness cannot be assessed for apparent motion, although it remains a factor in brushing-like motion.

It is also important to note that the study population in this experiment was limited, despite being comparable in size to prior studies. The sample primarily consisted of young adults. The narrow age range could limit the generalizability of the findings. Ageing is associated with a loss of peripheral afferents (Ekman et al. 2022) and also with an increase in perceived pleasantness of touch(Croy et al. 2019; Dione et al. 2023). Since these effects have not been investigated in relation to the perception of apparent motion, a sample of older participants could reveal different patterns of velocity-dependent pleasantness.

## Acknowledgements

Bengt Ragnemalm (Department of Biomedical and Clinical Sciences, Linköping University, Linköping, Sweden) provided essential technical and logistical support during the entire length of this project. The recruitment of participants was made possible by the help of Lotta Medling (Center for Social and Affective Neuroscience, Linköping University, Linköping, Sweden).

This research was supported by the Swedish Research Council (2020-01085, 2024-00381). Additionally, LJP was supported by the Medical Faculty of Linköping University, Linköping, Sweden.

## Data availability

Data and code used in this project is available through the projects Open Science Framework repository at https://osf.io/fj7xu/.

