## Supplementary Figure 1 for "Tactile illusion reveals central neural basis for touch pleasantness"

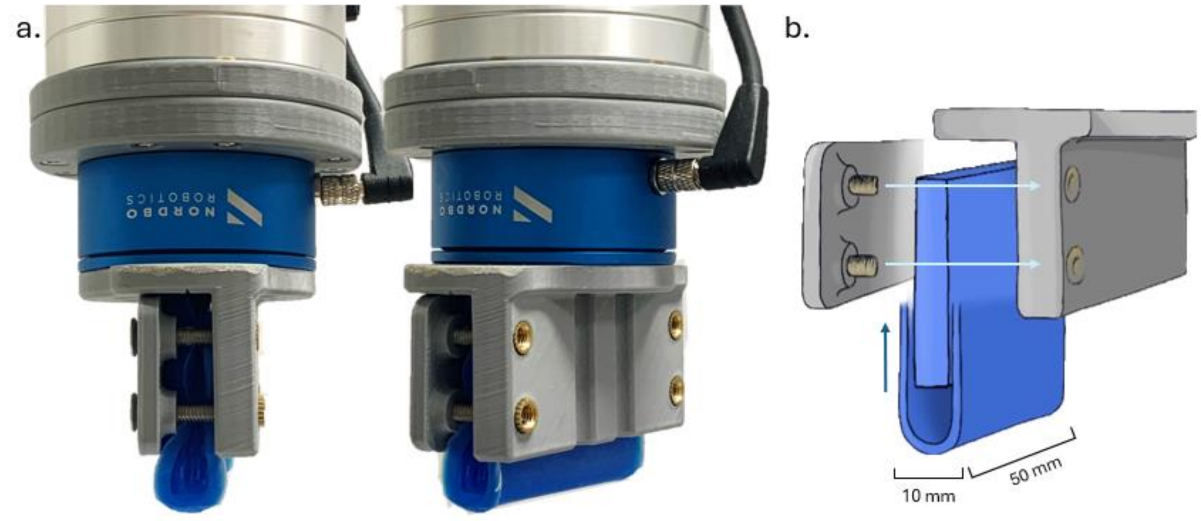


***Supplementary Figure 1*** **A.** Picture showing the silicone contactor, external force sensor and the end of the UR3e arm, from two different angles. **B.** Simple schematic breaking down the structure of the silicone contactor.
