## Supplementary Figure 3 for "Tactile illusion reveals central neural basis for touch pleasantness"

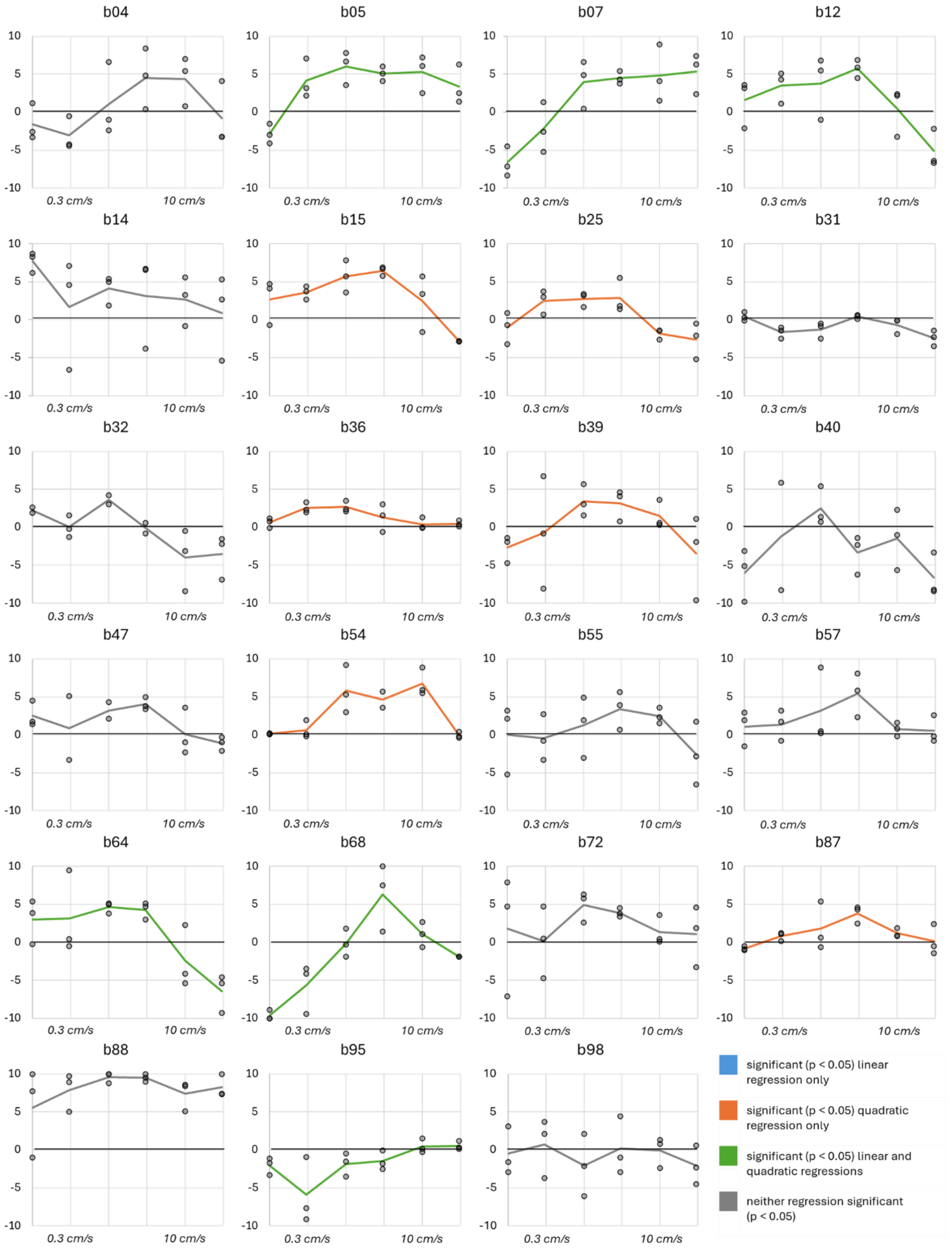


***Supplementary Figure 3*** Plots displaying pleasantness-ratings (y-axis) to brushing-like motion on the individual level. Velocity on log10-scale is shown on the x-axis, labelled with true velocities. Above each graph, the double-randomized participant ID is presented. Lines visualize the mean rating for each velocity respectively. The significance of the linear and quadratic regression can be interpreted from the colour of the line: blue for significant linear regression only, orange for significant quadratic regression only, green for significant linear and quadratic regression, and grey for neither regression reaching significance.
