## Supplementary Figure 4 for "Tactile illusion reveals central neural basis for touch pleasantness"

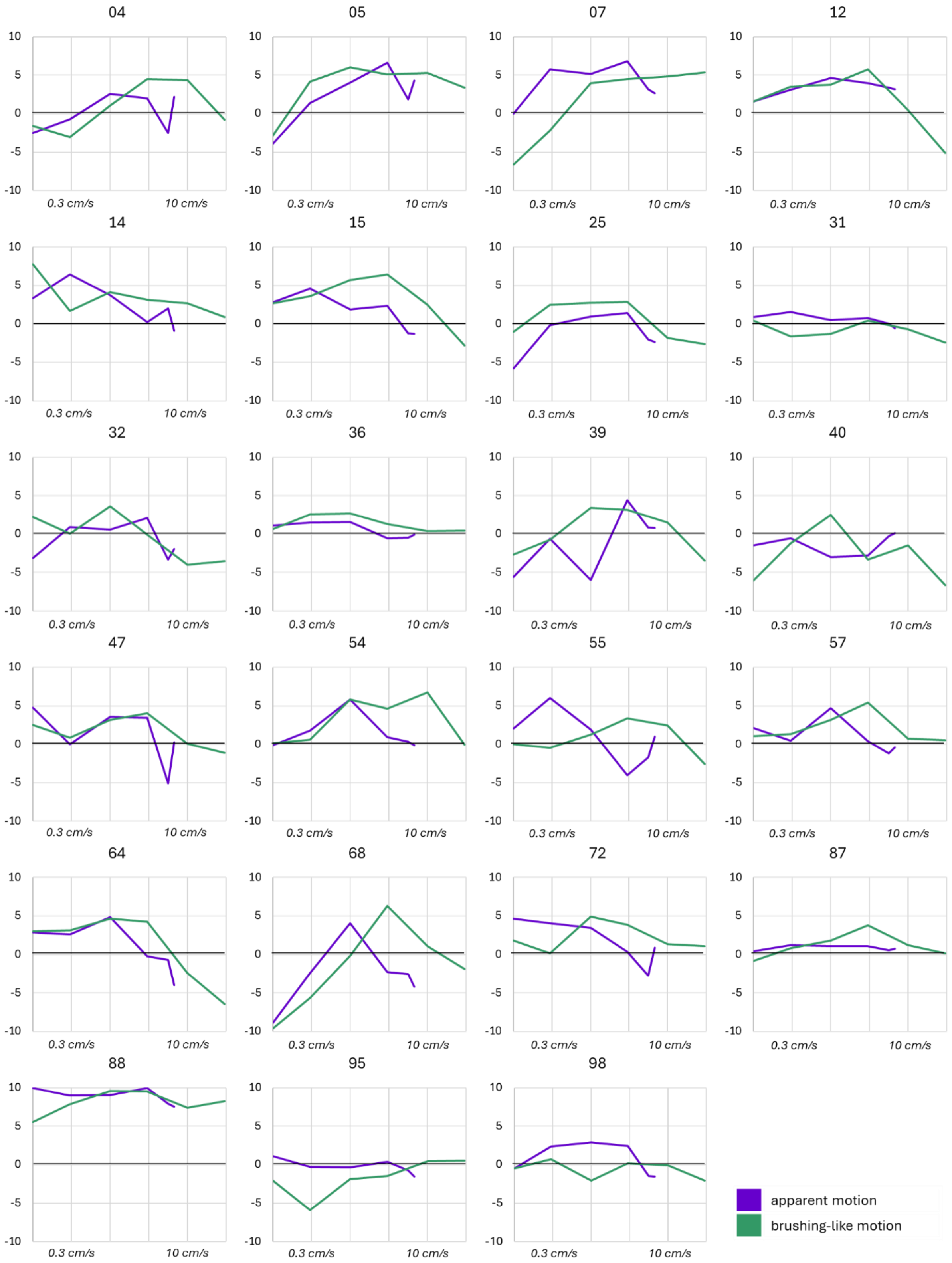


***Supplementary Figure 4*** Plots displaying pleasantness-ratings (y-axis) to apparent motion (purple) and brushing-like motion (green) on the individual level. Velocity on log10-scale is shown on the x-axis, labelled with true velocities. Above each graph, the double-randomized participant ID is presented.
