## Supplementary Figure 5 for "Tactile illusion reveals central neural basis for touch pleasantness"

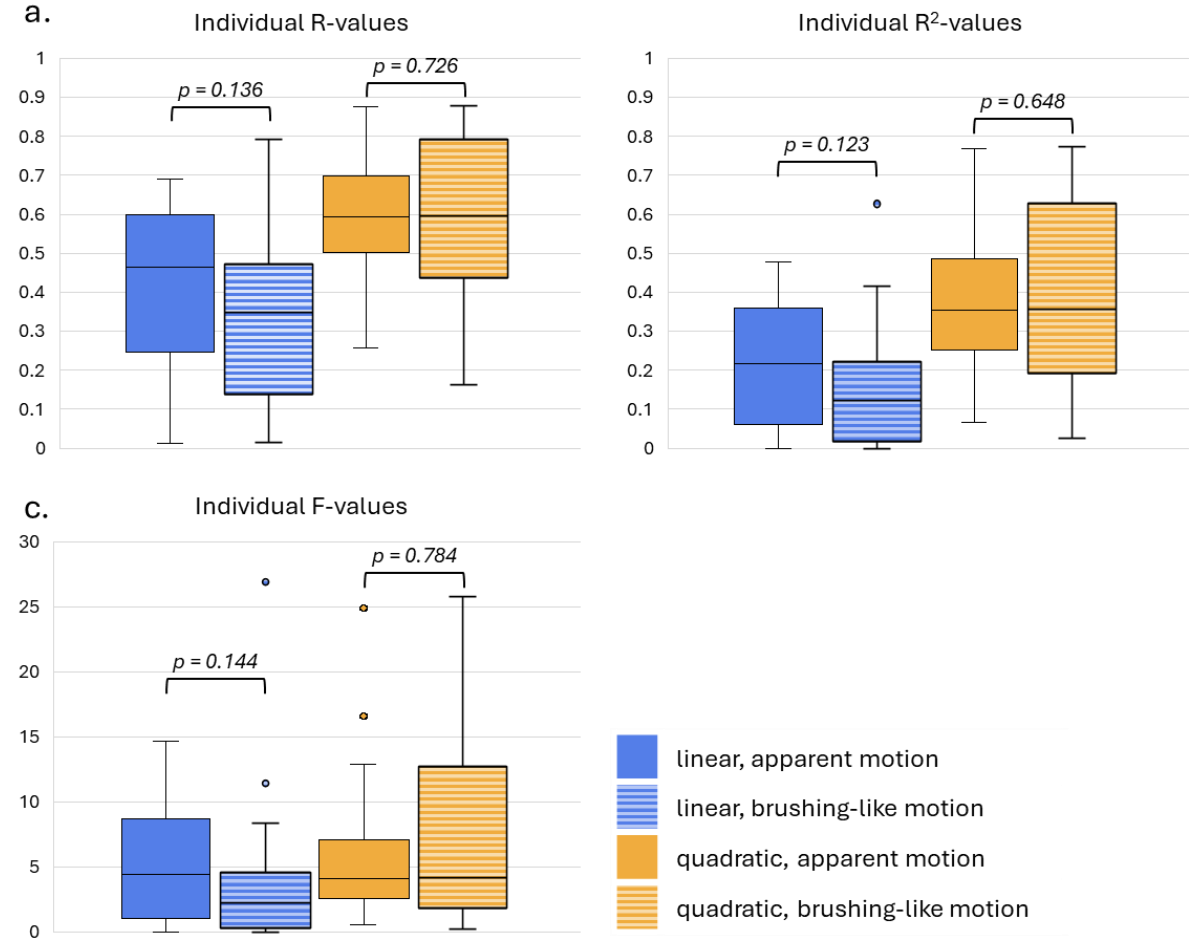


***Supplementary Figure 5*** Box plots showing the distribution of **a.** Individual R-values, **b.** Individual $R^2$-values and **c.** Individual F-values for the linear and quadratic regressions for both apparent motion and brushing-like motion. P-values comparing the motion conditions (Related-Samples Wilcoxon Signed Rank Test) for both regressions and the three values are displayed above the respective boxplots.
