## Supplementary Figure 6 for "Tactile illusion reveals central neural basis for touch pleasantness"

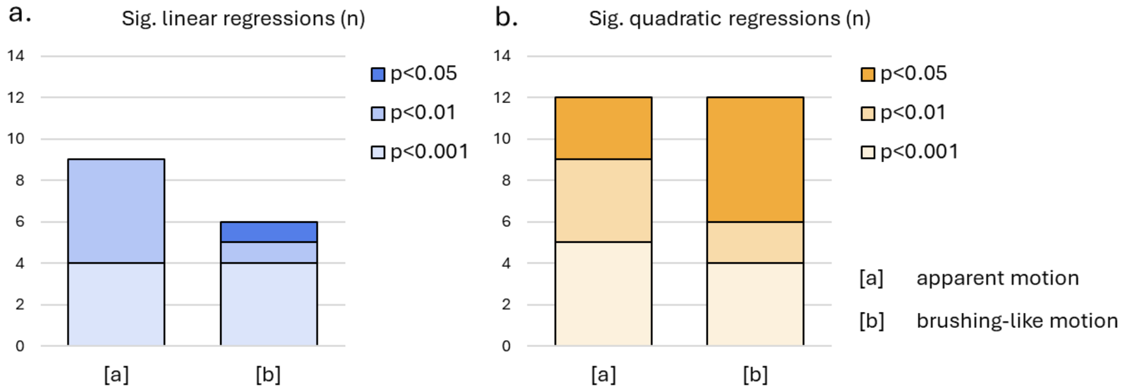


***Supplementary Figure 6*** Histograms showing the number of **a.** Significant linear and **b.** Significant quadratic regression for apparent motion [a] and brushing-like motion [b]. The shade of the stacked histogram indicates the significance level: deepest shade for p-values [0.01, 0.05), medium shade for [0.001, 0.01) and the lightest shade for p-values < 0.001.
