## Supplementary Figure 7 for "Tactile illusion reveals central neural basis for touch pleasantness"

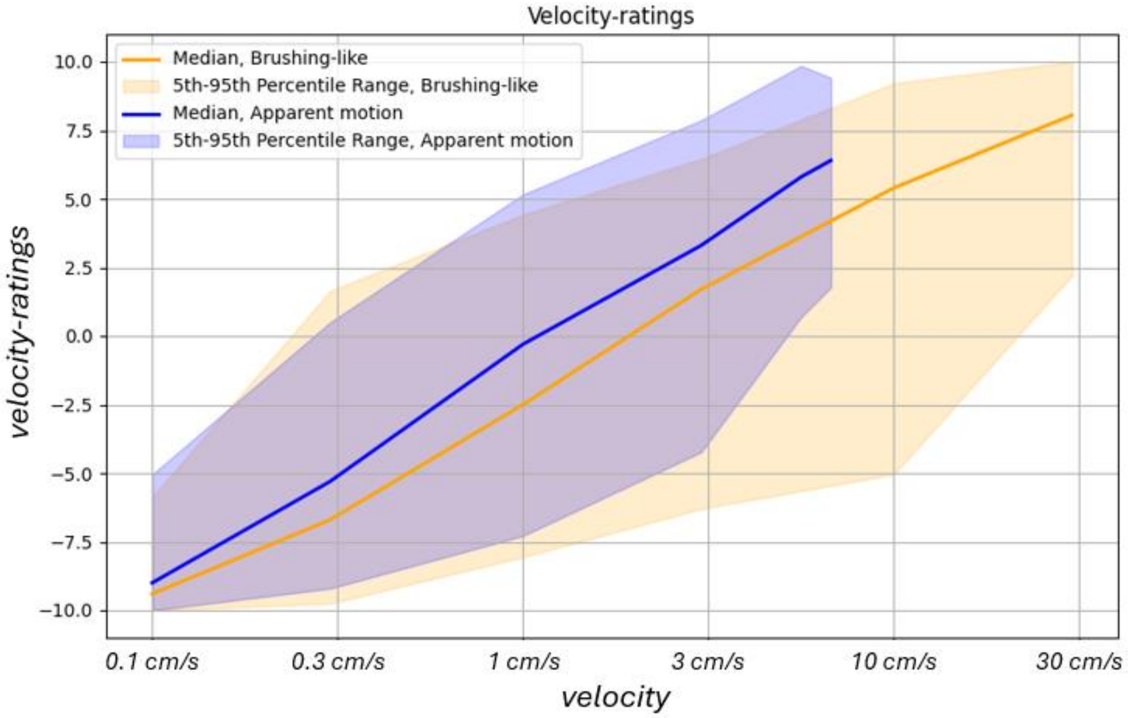


***Supplementary Figure 7*** Plot displaying the distribution of subjective velocity-ratings (y-axis) across the log10-transfromation of the global stimulus velocity (x-axis), labelled with the true velocities. Lines (blue – apparent motion, orange – brushing-like motion) present the median. The area between the 5th and the 95th percentiles is filled with corresponding colours.
