## Supplementary Table 1 for "Tactile illusion reveals central neural basis for touch pleasantness"

***Supplementary Table 1*** Coefficients for the linear and quadratic regressions, with pleasantness-ratings as the dependent variable and log10(velocity) and log10(velocity)2 as the predictors.

| stimulus | model | | Unstandardized B | Std. Error | Standardized Coefficients Beta | t | Sig. |
|---|---|---|---|---|---|---|---|
| apparent motion | Linear[1] | $\log_{10}$(velocity) | -0.522 | 0.278 | -0.093 | -1.878 | 0.061 |
| | Quadratic[2] | $\log_{10}(\text{velocity})^2$ | -2.979 | 0.559 | -1.036 | -5.327 | $1.03\times10^{-12}$ |
| | | $\log_{10}$(velocity) | 5.132 | 1.095 | 0.912 | 4.687 | $3.80\times10^{-6}$ |
| brushing-like motion | Linear[1] | $\log_{10}$(velocity) | 0.037 | 0.246 | 0.008 | 0.151 | 0.880 |
| | Quadratic[2] | $\log_{10}(\text{velocity})^2$ | -2.441 | 0.320 | -1.275 | -7.618 | $1.85\times10^{-13}$ |
| | | $\log_{10}$(velocity) | 6.088 | 0.827 | 1.232 | 7.362 | $1.66\times10^{-7}$ |

[1] Predictors: (Constant), $\log_{10}$(velocity)
[2] Predictors: (Constant), $\log_{10}$(velocity), $\log_{10}(\text{velocity})^2$
