## Supplementary Table 2 for "Tactile illusion reveals central neural basis for touch pleasantness"

***Supplementary Table 2*** Estimates of Fixed effects from the multilevel mixed effects analysis. Additionally, the F-values from type III Tests of Fixed Effects are displayed.

| Parameter | Estimate | 95% Confidence Interval | | Std. Error | t | F[c] | Sig. |
|---|---|---|---|---|---|---|---|
| | | Lower Bound | Upper Bound | | | | |
| Intercept | -0.815 | -1.633 | 0.004 | 0.417 | -1.954 | 0.423 | 0.0510 |
| [type=a] | 1.238 | 0.056 | 2.421 | 0.602 | 2.055 | 4.224 | 0.0402 |
| [type=b] | 0[b] | . | . | 0 | . | . | . |
| $\log_{10}$(velocity) | 6.088 | 4.529 | 7.648 | 0.794 | 7.664 | 65.646 | $5.13\times10^{-14}$ |
| $\log_{10}$(velocity)$^2$ | -2.441 | -3.045 | -1.837 | 0.308 | -7.930 | 68.245 | $7.20\times10^{-15}$ |
| [type=a] * $\log_{10}$(velocity) | -0.956 | -3.675 | 1.762 | 1.385 | -0.691 | 0.477 | 0.490 |
| [type=b] * $\log_{10}$(velocity) | 0[b] | . | . | 0 | . | . | . |
| [type=a] * $\log_{10}$(velocity)$^2$ | -0.539 | -1.827 | 0.749 | 0.656 | -0.821 | 0.674 | 0.412 |
| [type=b] * $\log_{10}$(velocity)$^2$ | 0[b] | . | . | 0 | . | . | . |

a. Dependent Variable: pleasantness, df = 815

b. This parameter is set to zero because it is redundant.

c. F-value for the type III Tests of Fixed Effects
