## Supplementary Table 3 for "Tactile illusion reveals central neural basis for touch pleasantness"

***Supplementary Table 3*** Related-Samples Wilcoxon Signed Rank Test comparing R-values, R2-values and F-values of the linear and quadratic models for the motion-conditions. Visual representation can be seen in Figure 5.

| Model | Pair | Sig.[1] | Median difference[2] | 95% Confidence Interval | |
|---|---|---|---|---|---|
| | | | | Lower | Upper |
| Linear | R-value apparent motion – R-value brushing-like motion | 0.136 | -0.090 | -0.219 | 0.036 |
| | R-square apparent motion – R-square brushing-like motion | 0.123 | -0.072 | -0.163 | 0.019 |
| | F-value apparent motion – F-value brushing-like motion | 0.144 | -1.763 | -4.705 | 0.626 |
| Quadratic | R-value apparent motion – R-value brushing-like motion | 0.726 | 0.008 | -0.081 | 0.088 |
| | R-square apparent motion – R-square brushing-like motion | 0.648 | 0.014 | -0.083 | 0.103 |
| | F-value apparent motion – F-value brushing-like motion | 0.784 | 0.330 | -1.398 | 2.364 |

[1] *Asymptotic significance displayed*
[2] *Estimation of the median of the difference, Related-Samples Hodges-Lehman Median Difference*
