## Supplementary Table 4 for "Tactile illusion reveals central neural basis for touch pleasantness"

***Supplementary Table 4*** Spearman's correlations, including the 95% Confidence Interval, between the subjective velocity-ratings and stimulus velocity for apparent motion and brushing-like motion.

| stimulus | | Spearman's rho | Sig. (2-tailed) | 95% Confidence Intervals (2-tailed)*,** | |
|---|---|---|---|---|---|
| | | | | Lower | Upper |
| apparent motion | velocity-rating – velocity | 0.858 | $5.46{\times}10^{-120}$ | 0.829 | 0.882 |
| brushing-like motion | velocity-rating – velocity | 0.870 | $1.45{\times}10^{-126}$ | 0.843 | 0.892 |

** Estimation is based on Fisher's r-to-z transformation.*

*** Estimation of standard error is based on the formula proposed by Fieller, Hartley, and Pearson.*
