## Supplementary Table 5 for "Tactile illusion reveals central neural basis for touch pleasantness"

***Supplementary Table 5*** Estimates of Fixed effects from the multilevel mixed effects analysis, including the parameters type, velocity-rating, and velocity-rating$^2$. Dependent variable pleasantness. Additionally, the F-values from type III Tests of Fixed Effects are displayed.

| Parameter | Estimate | 95% Confidence Interval | | Std. Error | t | F[c] | Sig. |
|---|---|---|---|---|---|---|---|
| | | Lower Bound | Upper Bound | | | | |
| Intercept | 2.584 | 1.959 | 3.212 | 0.320 | 8.076 | 103.222 | $2.41\times10^{-15}$ |
| [type=a] | -0.643 | -1.517 | 0.231 | 0.445 | -1.444 | 2.085 | 0.149 |
| [type=b] | 0[b] | . | . | 0 | . | . | . |
| velocity-rating | -0.013 | -0.072 | 0.047 | 0.030 | -0.425 | 3.301 | 0.671 |
| (velocity-rating)$^2$ | -0.032 | -0.043 | -0.020 | 0.006 | -5.292 | 36.081 | $1.56\times10^{-15}$ |
| [type=a] * velocity-rating | -0.055 | -0.143 | 0.032 | 0.045 | -1.241 | 1.539 | 0.215 |
| [type=b] * velocity-rating | 0[b] | . | . | 0 | . | . | . |
| [type=a] * (velocity-rating)$^2$ | 0.010 | -0.008 | 0.027 | 0.009 | 1.112 | 1.238 | 0.266 |
| [type=b] * (velocity-rating)$^2$ | 0[b] | . | . | 0 | . | . | . |

a. Dependent Variable: pleasantness, df = 815

b. This parameter is set to zero because it is redundant.

c. F-value for the type III Tests of Fixed Effects
